# Task-optimized neural networks reveal distinct contributions of specialized and broader visual learning to neural representations of face familiarity

**DOI:** 10.64898/2026.09.08.750266

**Authors:** Hamza Abdelhedi, Shahab Bakhtiari, Karim Jerbi

## Abstract

How neural activity across the ventral visual hierarchy supports face recognition is an open question. A long-standing debate asks whether face processing, particularly in fusiform cortex, relies on face-specific computations or representations shared with broader visual recognition. Here we combine source-resolved magnetoencephalography (MEG) with task-optimized neural networks as controlled computational models of visual experience. Rather than manipulating long-term expertise in human observers, we systematically vary learning objective on the model side—what the networks are trained to recognize—while holding architecture and loss function constant within model comparisons. We then ask which learned representational geometries best align with neural responses, where and when. We measured millisecond-resolved brain–model alignment across V1, lateral occipital cortex (LOC) and fusiform cortex while participants viewed familiar, unfamiliar and scrambled faces. The same stimuli were presented to seven CNN architectures trained for face-identity recognition (FR), object-category recognition (OR) or object categorization including a face category (Dual), alongside untrained controls. Familiarity produced a stage-dependent dissociation: in LOC, familiar faces showed earlier brain–model alignment than unfamiliar faces in the M170 range, an effect most consistent in models trained for face-recognition, whereas broader objectives produced more variable, architecture-dependent peak alignment latencies. In fusiform cortex they showed stronger alignment around the M200 range. This fusiform advantage was not uniquely associated with face-recognition training: dual- and object-trained models showed greater fusiform correspondence than face-trained models. Together, these findings support stage-dependent specialization, with training for face-identity recognition constraining intermediate-stage timing while later fusiform representations remain compatible with representational structure acquired through broader visual computations.

**Significance Statement:** Recognizing a familiar face feels immediate, yet it remains unclear which stages of visual processing are specifically shaped by learning individual identities. We combine millisecond-resolved MEG with task-optimized neural networks used as controllable models of visual experience, manipulating learning objective on the model side—that is, what the networks are trained to recognize and tracking when and where the resulting representations align with human brain activity. Familiarity advances representational alignment in lateral occipital cortex but strengthens later alignment in fusiform cortex. The earlier LOC effect was most consistent across architectures after face-identity recognition training, whereas the later fusiform effect also emerged under broader visual learning objectives. These findings support a stage-dependent account of face recognition that moves beyond a simple face-specific versus broader-visual-processing dichotomy.

## Introduction

How neural activity across the ventral visual hierarchy supports face recognition is a central question in visual neuroscience. A long-standing debate concerns the extent to which face processing depends on computations specialized for faces versus representational mechanisms shared with broader visual recognition (Kanwisher et al., 1997; Haxby et al., 2001; Gauthier and Tarr, 2002). This question has been especially prominent for fusiform cortex and the Fusiform Face Area (FFA), whose strong face selectivity has motivated domain-specific accounts, but whose responses have also been interpreted in relation to visual expertise and fine-grained individuation. Recent behavioral, neurophysiological, and computational work increasingly challenges a strict dichotomy between these views, showing that face-selective representations can coexist with broader dimensions of object representation and that functional specialization can emerge through task optimization (Bao et al., 2020; Dobs et al., 2022, 2023; Vinken et al., 2023). This reframes the debate from asking whether face processing relies on face-specific computations to asking where and when face-identity learning most strongly shapes representational dynamics along the visual hierarchy, and which stages remain compatible with representational structure acquired through broader visual learning.

Face familiarity provides a particularly informative window onto this question. Familiar and unfamiliar faces belong to the same visual category but differ profoundly in accumulated experience and access to stable identity representations (Gobbini and Haxby, 2007; Kramer et al., 2018; Wiese et al., 2024). They engage overlapping components of the distributed face-processing system (Haxby et al., 2000, 2001; Ramon et al., 2010; Duchaine and Yovel, 2015; Visconti di Oleggio Castello et al., 2017), yet where and when familiarity-related differences emerge remains debated (Wiese et al., 2024). Temporally, face perception unfolds over the first several hundred milliseconds after stimulus onset (Jeffreys, 1996; Liu et al., 2002; Vida et al., 2017). The face-sensitive M/N170 response, peaking around 150–170 ms (Liu et al., 2000; Halgren et al., 2000; Xu et al., 2005), has been associated with structural and configural processing of faces (Bentin et al., 1996; Eimer, 2000), with MEG source evidence localizing this response to the occipitotemporal cortex, including fusiform regions (Deffke et al., 2007). Later responses around 200 ms have been associated more consistently with identity-specific and familiarity-related processing (Halgren et al., 2000; Schweinberger et al., 2002; Gosling and Eimer, 2011; Wiese et al., 2024).

Evidence for familiarity effects across these temporal stages remains mixed, with electrophysiological work suggesting that familiarity can influence distinct stages of the visual processing hierarchy (Collins et al., 2018). Some studies report modulation in the M/N170 range (Caharel et al., 2002; Caharel and Rossion, 2021), whereas others find early responses comparatively insensitive to familiarity and place more reliable effects at later latencies (Bentin and Deouell, 2000; Schweinberger et al., 2002; Tanaka et al., 2006; Gosling and Eimer, 2011; Huang et al., 2017; Wiese et al., 2024). Time-resolved multivariate studies likewise reveal familiarity effects at multiple stages, including early enhancement of identity representations for familiar faces (Dobs et al., 2019) and later emergence of more image-tolerant identity representations (Ambrus et al., 2019; Ramon et al., 2015). Cross-experiment decoding further suggests that a general familiarity signature can emerge at later stages and generalize across different forms of familiarization (Dalski et al., 2022).

Spatially, lateral occipital cortex (LOC) is implicated in higher-level object processing and carries information relevant to facial identity (Grill-Spector et al., 2001; Vida et al., 2017), whereas fusiform cortex has been strongly implicated in face-selective and identity-sensitive processing (Kanwisher et al., 1997; Gauthier et al., 2000; Nasr and Tootell, 2012; Visconti di Oleggio Castello et al., 2017). Yet even the direction of familiarity effects in fusiform cortex has varied across studies, with reports of stronger responses for unfamiliar faces (Rossion et al., 2003) and others showing stronger responses for familiar faces (Weibert and Andrews, 2015). Together, this literature leaves unresolved whether familiarity primarily shifts the timing at which representations emerge, changes the strength with which they are expressed, or differentially affects these properties across successive stages of the ventral hierarchy.

A major challenge in addressing this question is that the learning history underlying long-term human face familiarity cannot readily be manipulated experimentally. We therefore use task-optimized neural networks as controlled models of visual experience, an approach that allows the consequences of optimizing systems for different recognition demands to be examined directly (Kanwisher et al., 2023). Rather than attempting to manipulate long-term expertise in human observers, we systematically vary the learning objective on the model side—that is, what the networks are trained to recognize—while using the same architecture and loss function across objectives. We then ask which learned representational geometries best align with human neural responses, where and when. Brain–model comparisons have shown that task-optimized networks can capture aspects of representational geometry in visual cortex (Yamins et al., 2014; Güçlü and van Gerven, 2015; Cadena et al., 2019; Kubilius et al., 2019; Grossman et al., 2019), while representational similarity analysis (RSA) provides a common framework for comparing model and neural representations without requiring unit-to-unit correspondence (Kriegeskorte et al., 2008). Importantly, comparing the same architecture across learning objectives holds architectural constraints constant, allowing effects associated with learning objective to be examined within a common architecture. Comparing multiple architectures, in turn, tests whether these effects are consistent across implementations or depend on particular architectural constraints (Storrs et al., 2021). Direct comparisons of networks optimized for face-identity versus object recognition show that learning objective can shape the representations used for face recognition (Abudarham et al., 2021), while broader computational work demonstrates that learning objective can shape the emergence of category-selective and functionally specialized representations in neural networks (Dobs et al., 2022, 2023; Prince et al., 2024).

Here, we combine source-resolved magnetoencephalography (MEG) with time-resolved RSA to quantify brain–model representational alignment at millisecond resolution across V1, LOC, and fusiform cortex. MEG provides the temporal precision required to track rapidly evolving representational dynamics, while source reconstruction enables these dynamics to be examined regionally (Cichy et al., 2017; Kietzmann et al., 2019; Dehgan et al., 2025). Human MEG responses were measured for familiar, unfamiliar, and scrambled faces, and the same stimuli were presented to seven convolutional neural network architectures originally developed for face or object recognition. Each architecture was trained under three learning objectives—face-identity recognition (FR), object-category recognition (OR), or object categorization including a face category (Dual)—with an untrained counterpart serving as a control. Scrambled faces provided a control for correspondence arising from visual responses in the absence of intact face structure, whereas untrained networks tested the contribution of architecture in the absence of learning.

We address two complementary questions. First, where along the ventral visual hierarchy does familiarity alter brain–model representational alignment, and is this expressed through changes in timing, strength, or both? Second, which familiarity-sensitive alignment dynamics are most strongly shaped by the face-identity recognition objective, and which also emerge under broader visual learning objectives? By resolving brain–model correspondence jointly across region, time, architecture, stimulus familiarity, and learning objective, we aim to move beyond a simple face-specific-versus-domain-general dichotomy and clarify how face-identity and broader visual learning shape representations across successive stages of human face processing.

## Results

To determine where and when face familiarity shapes visual representations, we compared source-localized MEG responses to familiar, unfamiliar, and scrambled faces with representations learned by task-optimized neural networks (Fig. 1). MEG activity was resolved across 448 cortical parcels from which time-resolved representational dissimilarity matrices (RDMs) were computed separately for each stimulus condition. The same images were presented to seven CNN architectures—FaceNet, SphereFace, CORnet-S, ResNet50, VGG16, Inception, and MobileNetV2—trained under three learning objectives: face-identity recognition (FR), object-category recognition (OR), or object categorization including a face category (Dual). Untrained counterparts were included to isolate the contribution of learning from architecture alone.

**Figure 1.**
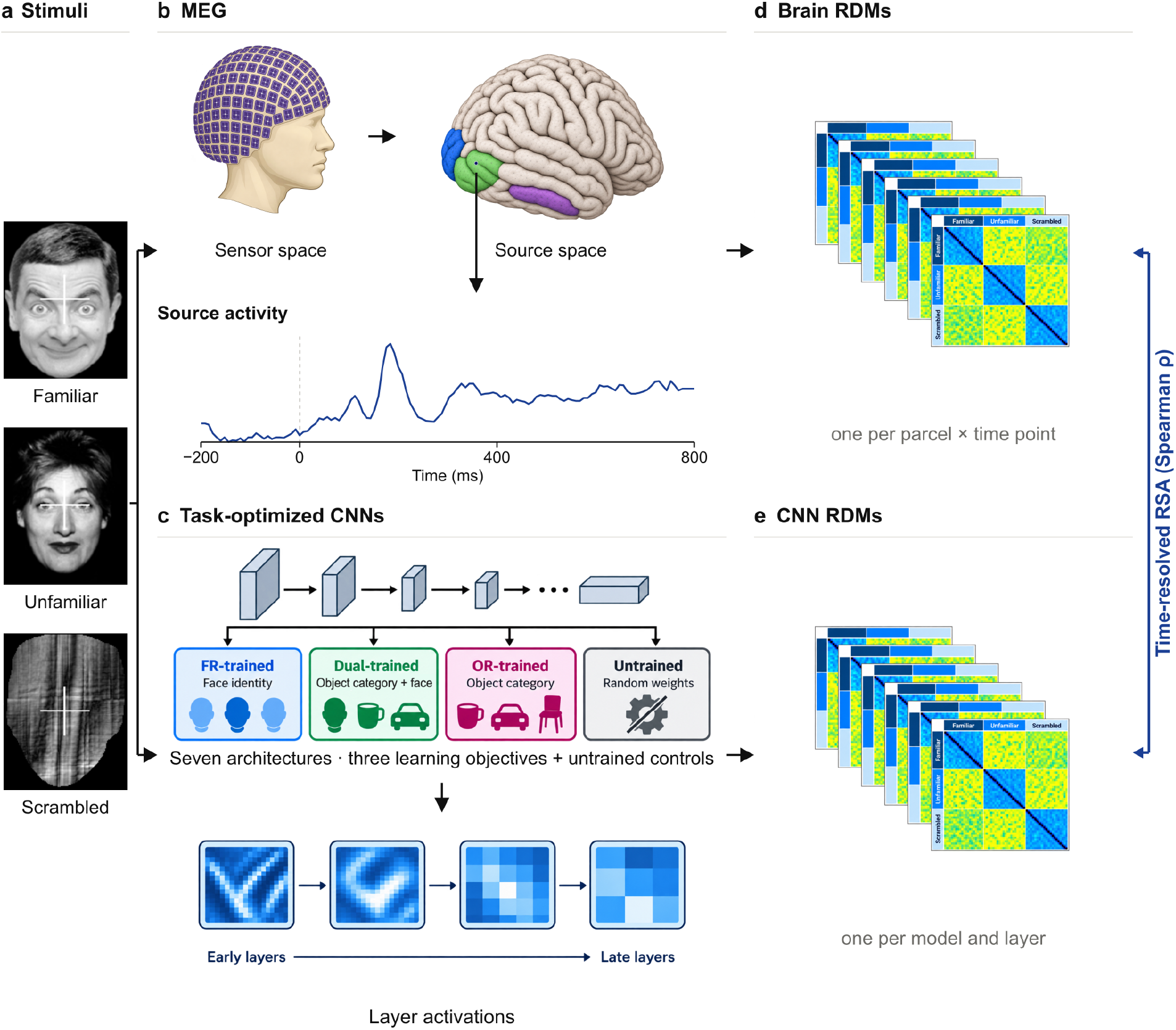
Study design. **a, Stimuli**. Participants viewed familiar, unfamiliar, and scrambled faces. **b, MEG**. Sensor-space MEG data were source-localized to cortical regions including V1, LOC, and fusiform cortex, yielding time-resolved source-level activity. **c, Task-optimized CNNs**. Seven CNN architectures were examined under three learning objectives: face-identity recognition (FR), object-category recognition (OR), and object categorization including a face category (Dual), together with untrained controls. Activations were extracted across network layers. **d, Brain RDMs**. For each cortical parcel and time point, a representational dissimilarity matrix (RDM) captured pairwise dissimilarities among familiar, unfamiliar, and scrambled stimuli. **e, CNN RDMs**. For each model and layer, an RDM captured pairwise dissimilarities among the same stimuli. Condition-specific sub-RDMs were extracted from the full RDMs and compared using time-resolved representational similarity analysis (RSA; Spearman correlation) at each MEG time point × CNN layer combination. Peak-strength analyses used the maximum over this grid, whereas time-resolved analyses visualized the layer yielding maximal similarity for the relevant condition. Model-level analyses used subject-averaged MEG RDMs, whereas subject-level robustness analyses used individual-participant MEG RDMs. Similarity scores were normalized by the upper noise ceiling.

Brain–model alignment was quantified using representational similarity analysis (RSA) by comparing each time-resolved MEG RDM with layer-resolved model RDMs for the corresponding stimulus condition (Fig. 1). Similarity values were normalized by the MEG noise ceiling. Unless otherwise stated, we report model-level analyses based on subject-averaged MEG RDMs and focused on three literature-guided right-hemisphere regions spanning early and ventral visual cortex: V1, lateral occipital cortex (LOC), and fusiform cortex. Subject-level analyses were used as complementary robustness tests.

### Familiarity selectively strengthens fusiform brain–model alignment

We first asked whether face familiarity alters the strength of brain–model correspondence in LOC and fusiform cortex. For each architecture, learning objective, and stimulus condition, we quantified the maximum MEG–CNN RSA similarity across the time *×* layer space (Fig. 2a,b). This measure captures the strongest representational correspondence for each model–condition pair, irrespective of when it occurs or which model layer contributes most strongly; temporal differences are examined separately below. Pairwise comparisons of peak similarity were performed across the seven matched architectures.

**Figure 2.**
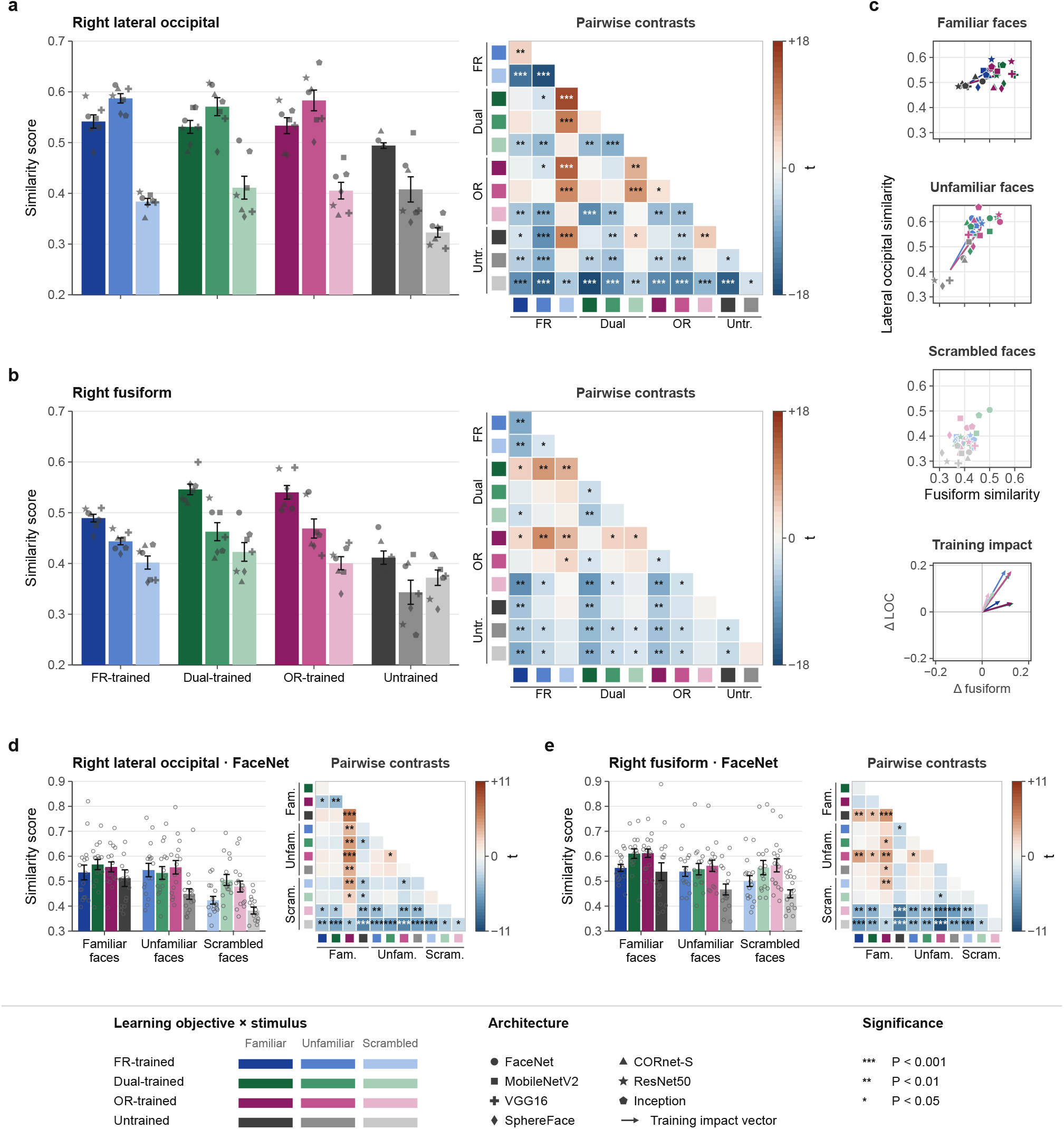
Peak brain–CNN representational similarity in right lateral occipital and fusiform cortex. (**a**) Right lateral occipital cortex (LOC): maximum MEG–CNN RSA similarity across time and network layers for familiar, unfamiliar, and scrambled faces. Similarity was computed from subject-averaged MEG RDMs (*n*=16) for seven CNN architectures (*n*=7; each point represents one architecture) under FR-trained, Dual-trained, OR-trained, and untrained conditions. The adjacent heat map shows pairwise contrasts between learning objective *×* stimulus conditions, expressed as *t* values from two-sided paired tests across architectures. (**b**) Same analysis for right fusiform cortex. (**c**) Cross-region correspondence between LOC and fusiform peak similarity, shown separately for familiar, unfamiliar, and scrambled faces. Each point represents one architecture. The training-impact panel shows the change from each architecture’s untrained baseline to its trained counterpart, illustrating how training shifts peak correspondence across the two regions. (**d**) Subject-level FaceNet analysis in right LOC. Bars show peak similarity for familiar, unfamiliar, and scrambled faces across participants (*n*=16), with individual participants shown as points. The adjacent heat map summarizes pairwise contrasts with *t* values from two-sided paired tests across participants. (**e**) Same subject-level analysis for right fusiform cortex. Similarity scores were normalized by the upper noise ceiling.

In LOC, peak alignment did not differ significantly across learning objectives (Fig. 2a), nor was it enhanced by familiarity. Instead, unfamiliar faces showed slightly greater peak similarity than familiar faces, an effect that was significant for FR-trained models (*t* = 4.2, *p <* 0.01), weaker for OR-trained models (*t* = 2.9, *p <* 0.05), and not significant for Dual-trained models (*t* = 2.1, *p >* 0.05). Thus, LOC showed no familiarity advantage in peak correspondence.

A markedly different pattern emerged in fusiform cortex (Fig. 2b). Across all three learning objectives, familiar faces showed stronger peak alignment than unfamiliar faces (FR: *t* = −8.5, *p <* 0.01; Dual: *t* = −3.7, *p <* 0.05; OR: *t* = −2.9, *p <* 0.05). Moreover, for familiar faces, Dual- and OR-trained networks showed greater fusiform correspondence than FR-trained networks (FR *<* OR: |*t*| = 3.5, *p <* 0.05; FR *<* dual: |*t*| = 4.4, *p <* 0.05). Thus, the fusiform familiarity advantage was robust across learning objectives rather than specific to face-recognition training.

The joint LOC–fusiform representation provided a complementary visualization of this regional dissociation (Fig. 2c). Training shifted models toward stronger alignment in both regions, with architectures showing different spatial distributions across learning objective. Familiarity, however, affected the two regions differently: familiar-face representations shifted toward greater fusiform correspondence, whereas LOC showed little or an opposite change in peak strength. FR-trained architectures appeared more tightly grouped than dual- and OR-trained models, indicating greater consistency across architectures under this objective.

### Control and robustness analyses support the specificity of brain–model alignment

Several complementary analyses tested whether these effects could be explained by generic visual responses, architecture alone, or spatially nonspecific brain–model correspondence. In V1, similarity was lower overall, and scrambled faces often produced comparatively strong correspondence (Supplementary Fig. S1a). By contrast, familiar and unfamiliar faces generally aligned more strongly than scrambled stimuli in LOC and fusiform cortex. Trained networks also consistently outperformed their untrained counterparts, indicating that correspondence depended on learned representational structure rather than architecture alone. We further examined two control regions—the entorhinal cortex and caudal anterior cingulate—where similarity was low, learning objectives showed no systematic clustering, and training-impact vectors were short and inconsistent (Supplementary Fig. S1c). Together, these controls indicate that the MEG–CNN correspondence observed along the ventral visual stream was not a ubiquitous feature of cortical activity.

Subject-level analyses provided an additional robustness test (Fig. 2d,e). In LOC, FaceNet-based similarity varied substantially across participants and showed no reliable familiarity effect (Fig. 2d). In fusiform cortex, however, familiar faces again showed stronger alignment than unfamiliar faces, reaching significance for Dual-trained FaceNet and appearing less variable across participants (Fig. 2e). Supplementary analyses with CORnet-S and ResNet50 showed a similar regional pattern, although correspondence was weaker and reduced statistical significance relative to FaceNet. Frequency-resolved analyses further showed the strongest correspondence in gamma and high-gamma activity (Supplementary Fig. S2).

### Face familiarity yields earlier brain–model alignment in LOC

Having found that familiarity increased the strength of brain–model correspondence in fusiform cortex but not in LOC, we next asked whether it also altered the timing of representational alignment. We therefore examined time-resolved MEG–CNN similarity across V1, LOC, and fusiform cortex using FR-trained FaceNet for detailed time-resolved analyses (Fig. 3a–i), before comparing peak alignment latencies across all seven FR-trained architectures (Fig. 3j).

**Figure 3.**
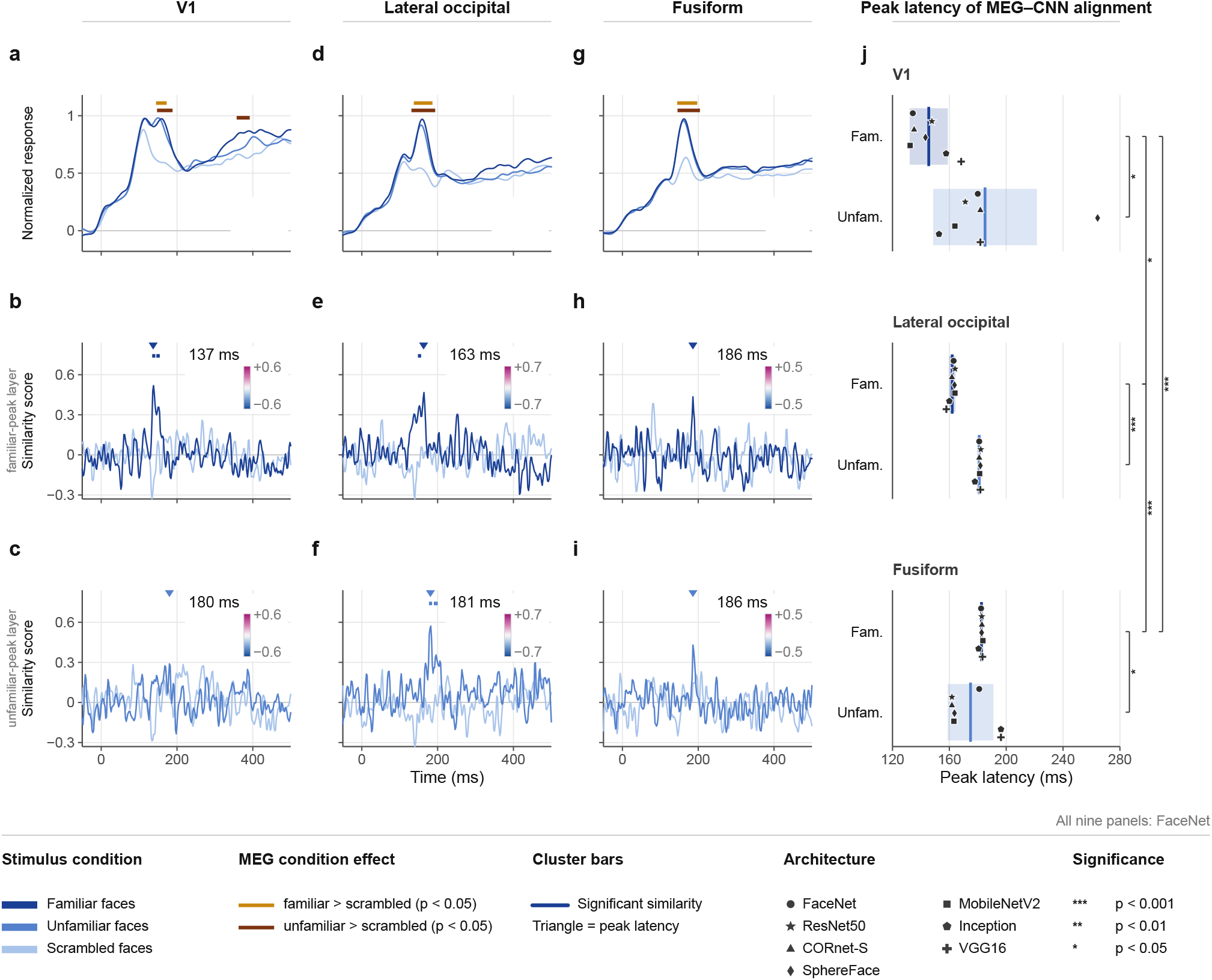
Spatiotemporal dynamics of MEG–CNN similarity for FR-trained models across V1, lateral occipital cortex, and fusiform cortex. (**a**,**d**,**g**) Source-level MEG responses to familiar, unfamiliar, and scrambled faces in V1 (a), lateral occipital cortex (LOC; d), and fusiform cortex (g). Yellow and orange bars indicate significant familiar *>* scrambled and unfamiliar *>* scrambled effects, respectively (cluster-based permutation tests, *p <* 0.05). (**b**,**e**,**h**) Time-resolved MEG–FaceNet RSA similarity in V1 (b), LOC (e), and fusiform cortex (h), evaluated at the FaceNet layer yielding maximal similarity for familiar faces. Triangles mark peak similarity latencies, and inset cortical maps show the spatial distribution of similarity at the corresponding peak latency. (**c**,**f**,**i**) Corresponding time-resolved analyses evaluated at the FaceNet layer yielding maximal similarity for unfamiliar faces in V1 (c), LOC (f), and fusiform cortex (i). (**j**) Peak latency of MEG–CNN alignment for familiar and unfamiliar faces across V1, LOC, and fusiform cortex, computed for each of the seven FR-trained architectures within the 50–400 ms search window. Each point represents one architecture; bars indicate mean *±* SD. Brackets denote significant paired comparisons across regions and stimulus conditions.

In V1, source-level responses showed two early deflections between approximately 110 and 150 ms, with the later component more pronounced for faces than for scrambled stimuli (Fig. 3a). Brain–model similarity for familiar faces increased sharply around 130— 140 ms and formed a significant cluster peaking at 137 ms, whereas similarity for unfamiliar faces did not form a significant cluster in the FaceNet analysis (Fig. 3b). Scrambled faces elicited comparable early MEG responses but negligible model alignment, showing that measurable evoked activity alone did not guarantee substantial representational correspondence.

The clearest familiarity-related temporal effect emerged in LOC. Source-level activity showed a prominent face-related response in the 150–180 ms range, consistent with the M170 stage of face processing (Fig. 3d). Brain–model alignment for familiar faces formed a significant cluster encompassing approximately 140–170 ms and peaked at 163 ms, whereas unfamiliar faces showed a later significant cluster encompassing approximately 180–210 ms, peaking at 181 ms (Fig. 3e,f). Peak alignment strengths were comparable between the two conditions. Thus, within FaceNet, familiarity primarily altered the latency of LOC alignment: familiar faces reached peak correspondence approximately 18 ms earlier than unfamiliar faces without a corresponding increase in peak strength.

Fusiform cortex showed a different temporal profile. Both familiar and unfamiliar faces elicited robust source-level responses relative to scrambled controls (Fig. 3g), but the time-resolved brain–model similarity traces did not show the clear familiarity-related temporal separation observed in LOC. For FaceNet, both conditions reached their descriptive maximum at approximately 186 ms (Fig. 3h,i). Thus, although fusiform correspondence reached its maximum later in the M200 range, familiarity was not associated with a similarly robust advance in peak latency. This complements the preceding peak-strength analysis, in which the principal fusiform familiarity effect was stronger alignment for familiar than unfamiliar faces.

We next asked whether the earlier LOC alignment for familiar faces was also evident across the other architectures. Across the seven FR-trained architectures, familiar-face peak latencies progressed from V1 (143.7 *±* 11.4 ms) to LOC (161.9 *±* 2.2 ms) to fusiform cortex (185.4 *±* 1.4 ms), with significant pairwise differences between regions (Fig. 3j). Although familiar–unfamiliar peak-latency differences reached significance in all three ROIs, their stability across architectures varied markedly by regions. The LOC effect was particularly consistent across architectures (*p* = 5.14 *×* 10^−7^), with unfamiliar-face peaks systematically occurring later than familiar-face peaks. By contrast, unfamiliar-face peak estimates in V1 and fusiform cortex showed substantially greater between-model dispersion, resulting in greater architecture dependence of the corresponding latency differences.

Frequency-resolved analyses provided complementary context for the broadband findings (Supplementary Fig. S3). Familiarity-related temporal differences were most evident in LOC, particularly in theta and alpha activity, whereas fusiform similarity was generally weaker and showed less consistent temporal structure across bands. Scrambled stimuli again showed minimal RSA correspondence despite measurable band-limited responses.

Together, these analyses identify earlier LOC brain–model alignment as the most consistent temporal signature of face familiarity. Across successive visual stages, the familiarity difference was expressed primarily in alignment timing in LOC, whereas in fusiform cortex it was expressed predominantly in alignment strength.

### Face-identity recognition training yields the most consistent familiarity-related advance in LOC peak alignment

Having identified earlier LOC alignment as the clearest temporal signature of face familiarity, we next asked whether this effect depended on what the models had learned to recognize. We therefore compared networks trained for face-identity recognition, object-category recognition, or object categorization including a face category. We first examined these effects within FaceNet and then assessed their consistency across all seven architectures (Fig. 4).

**Figure 4.**
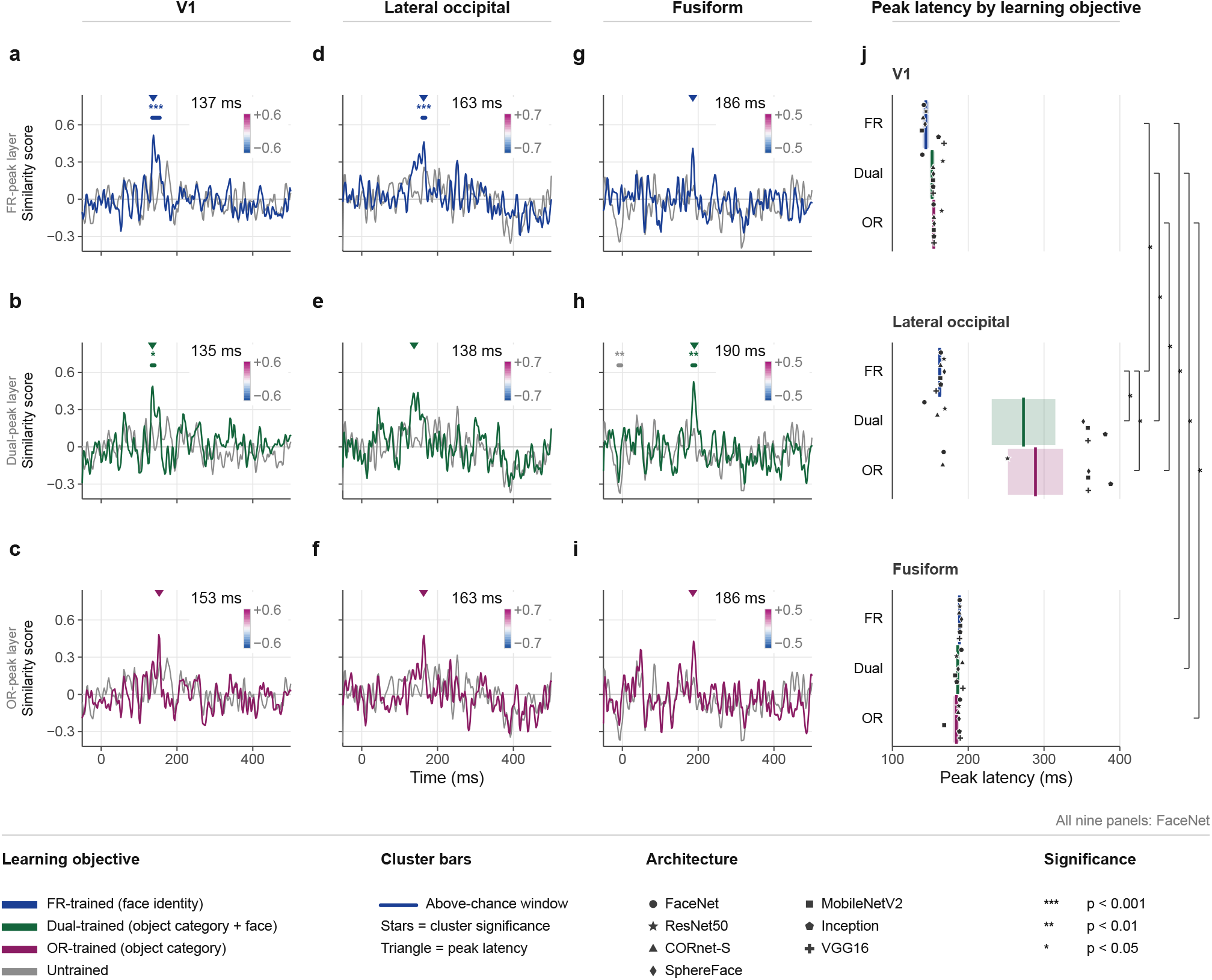
Learning objective shapes the spatiotemporal dynamics of MEG–CNN similarity for familiar faces across V1, lateral occipital cortex, and fusiform cortex. (**a**,**d**,**g**) Time-resolved MEG–FaceNet RSA similarity for familiar faces under the FR-trained objective in V1 (a), lateral occipital cortex (LOC; d), and fusiform cortex (g). Similarity is shown at the CNN layer yielding maximal familiar-face similarity for the FR-trained model; gray traces show the corresponding untrained model. Colored horizontal bars indicate significant above-chance similarity clusters, with stars denoting significance level, and triangles marking peak similarity latency. Inset cortical maps show the spatial distribution of similarity at the corresponding peak latency. (**b**,**e**,**h**) Corresponding analyses for the Dual-trained objective in V1 (b), LOC (e), and fusiform cortex (h), evaluated at the layer yielding maximal familiar-face similarity for the Dual-trained model. (**c**,**f**,**i**) Corresponding analyses for the OR-trained objective in V1 (c), LOC (f), and fusiform cortex (i), evaluated at the layer yielding maximal familiar-face similarity for the OR-trained model. (**j**) Peak latency of MEG–CNN alignment for familiar faces across V1, LOC, and fusiform cortex, shown separately for FR-trained, Dual-trained, and OR-trained models. Peak latency was defined as the time of maximal similarity within the 50–400 ms search window. Each point represents one of seven CNN architectures (FaceNet, ResNet50, CORnet-S, MobileNetV2, SphereFace, Inception, and VGG16); bars indicate mean *±* SD. Brackets denote significant paired comparisons across regions and learning objectives.

In V1, familiar-face alignment remained early under all three learning objectives, with FaceNet peaks at 137 ms for FR training, 135 ms for Dual training, and 153 ms for OR training (Fig. 4a-c). Thus, changing the learning objective produced only modest differences in the timing of early visual correspondence.

Learning objective had a much stronger effect in LOC (Fig. 4d-f). FR-trained FaceNet peaked at 163 ms, whereas Dual-trained FaceNet peaked even earlier, at 138 ms, and exhibited a more prolonged significant similarity cluster. OR-trained FaceNet peaked near the FR-trained latency but did not show a significant similarity cluster. Thus, within FaceNet alone, early LOC alignment was not exclusive to FR objective; an early peak was also observed under Dual and OR objectives, with the Dual model peaking earliest.

The cross-architecture analysis revealed a different pattern (Fig. 4j). Under the FR objective, familiar-face LOC peaks were tightly concentrated in the early M170 range across architectures (161.9 *±* 2.2 ms). Under Dual and OR objectives, peak latencies became markedly more dispersed (Dual: 272.9 *±* 111.1 ms; OR: 288.2 *±* 96.1 ms). Mean peak latencies differed significantly from FR training (*p* = 0.0395 and *p* = 0.0136, respectively). Importantly, these later group means did not reflect a uniform delay across architectures. Instead, architecture-level peaks separated into early and late groups: some architectures retained an M170-range peak, whereas others shifted toward much later peaks around 350 ms. The key effect of FR training was therefore not to create an early LOC solution that broader objectives could not achieve, but to make that early solution far more consistent across architectures.

This interpretation becomes especially clear when familiar and unfamiliar faces are considered together. For unfamiliar faces, LOC peak alignment converged near 180 ms across all three learning objectives, with little between-objective variation (Supplementary Fig. S4). Thus, the earlier LOC peak for familiar than unfamiliar faces was most consistently expressed in models trained for face recognition, whereas broader objectives produced more architecture-dependent familiar-face peak latencies. In other words, face-identity learning appears to constrain when familiar-face representations converge in LOC, while broader visual learning permits multiple temporal solutions.

### Fusiform familiarity effects extend beyond face-identity recognition training

We next asked whether the fusiform familiarity effect in the M200 range showed the same relationship to learning objective. Here, the pattern was notably different. In FaceNet, familiar-face alignment peaked at similar latencies under all three objectives—186 ms for FR training, 190 ms for dual training, and 186 ms for OR training (Fig. 4g-i). Despite their similar peak latencies, the temporal profiles differed in strength and duration: the Dual-trained model showed the clearest and most sustained significant fusiform correspondence, whereas the FR- and OR-trained variants showed briefer or non-significant peaks.

This relative stability was also evident across architectures. Unlike LOC, where broader learning objectives were associated with substantially greater cross-architecture dispersion in peak latency, fusiform peaks remained concentrated near the M200 range across learning objectives (FR: 185.4 *±* 1.4 ms; Dual: 186.4*±* 3.0 ms; OR: 180.7 *±* 12.8 ms; Fig. 4j). Learning objective therefore had a much stronger effect on the temporal stability of LOC alignment than on the timing of familiar-face fusiform correspondence.

More importantly, the strength of the fusiform familiarity effect was not specific to FR training. As shown in Fig. 2b, familiar faces produced stronger fusiform alignment than unfamiliar faces under all three learning objectives. Moreover, for familiar faces, Dual- and OR-trained models showed greater peak fusiform correspondence than FR-trained models. Thus, the familiar-face fusiform signature persisted—and could even be enhanced—when models were trained on broader visual objectives.

Taken together, Figures 2 and 4 reveal a clear stage-dependent difference in how learning objective shapes familiar-face representations. FR training most strongly constrains the timing of intermediate LOC alignment, whereas later fusiform familiarity effects are not uniquely dependent on face-identity training and remain compatible with representations acquired through broader visual learning. This dissociation argues against a single, uniformly face-specific processing regime and instead points to different computational constraints at successive stages of the ventral visual pathway.

## Discussion

The findings show that face familiarity affects neural representations differently across stages of the ventral visual hierarchy. In lateral occipital cortex (LOC), brain–model alignment peaked earlier for familiar than unfamiliar faces. In fusiform cortex, familiarity was primarily associated with stronger alignment, with less consistent evidence for a change in when that alignment peaked. Systematic comparison of model learning objective further showed that these signatures differed in how strongly they depended on face-identity learning. The earlier LOC alignment was expressed most consistently across architectures after face-identity recognition training, whereas the fusiform familiarity advantage emerged across all three learning objectives and was even stronger at peak for Dual- and OR-trained models than for FR-trained models. Together, these results support distinct roles for learning across successive stages of face processing: face-identity learning appears to constrain the temporal organization of intermediate LOC representations, while later fusiform familiarity representations remain substantially compatible with representational structure acquired through broader visual learning.

### Familiarity affects timing and strength at different stages

A central implication of our findings is that familiarity alters different properties of neural representations at successive stages of the ventral visual hierarchy. The progression from V1 to LOC to fusiform cortex broadly followed the canonical sequence of early visual, M170-range, and later M200-range responses described in previous electrophysiological work (Halgren et al., 2000; Liu et al., 2000). Within this sequence, however, familiarity was expressed differently across regions. In LOC, its clearest effect was temporal: familiar faces reached maximal brain–model alignment earlier than unfamiliar faces, with little corresponding familiarity advantage in peak strength. In fusiform cortex, by contrast, familiarity was expressed more reliably through stronger alignment than through a systematic shift in alignement peak latency. Thus, familiarity did not alter ventral-stream representations uniformly, but affected when representational alignment became maximal at an intermediate stage and how strongly it was expressed at a later stage.

This distinction offers one way to reconcile mixed reports of familiarity effects around the M/N170. Some studies have reported familiarity-related modulation in this range (Caharel et al., 2002; Caharel and Rossion, 2021), whereas others have found early responses comparatively insensitive to familiarity and located more robust effects at later stages (Bentin and Deouell, 2000; Schweinberger et al., 2002; Gosling and Eimer, 2011; Wiese et al., 2024). Our results suggest that, at intermediate stages of visual processing, face familiarity is reflected in earlier peaks of brain–model representational alignment. If familiarity primarily shifts the latency of peak brain–model alignment, analyses focused on neural response amplitude may miss this early timing difference between familiar and unfamiliar faces. Complementing these early timing differences, cross-experiment EEG decoding has identified a later familiarity signal, predominantly between 270 and 630 ms, that generalizes across participants, stimuli, and different forms of familiarization (Dalski et al., 2022).

The interpretation of this latency effect nevertheless requires precision. A peak in MEG–CNN RSA similarity identifies the time at which neural representational geometry most closely corresponds to a representation learned by the model. An earlier peak therefore indicates earlier maximal correspondence but does not by itself imply a larger neural response, faster neural computation, or faster behavioral recognition. Within these limits, the earlier LOC peak for familiar faces is consistent with earlier emergence or stabilization of familiarity-sensitive representational structure (Halgren et al., 2000; Liu et al., 2000; Cichy et al., 2017; Dobs et al., 2019). Conversely, the later peaks for unfamiliar faces may reflect additional feature integration or disambiguation before a comparable representational geometry becomes maximally expressed (Schweinberger et al., 2002; Gosling and Eimer, 2011; Caharel and Rossion, 2021). However, the computations responsible for this delay cannot be determined from representational timing alone.

The fusiform result points to a complementary form of familiarity sensitivity. Stronger correspondence for familiar than unfamiliar faces is consistent with accounts linking fusiform cortex to identity-sensitive and person-specific representations (Gobbini and Haxby, 2007; Weibert and Andrews, 2015; Wiese et al., 2024) and with MEG-RSA evidence for enhanced familiar-identity representations (Dobs et al., 2019). Importantly, representational correspondence and univariate response amplitude need not vary in parallel. This distinction may help contextualize apparently opposing reports of larger fusiform responses for unfamiliar faces (Rossion et al., 2003) and familiar faces (Weibert and Andrews, 2015): variability in response magnitude does not preclude a familiarity effect in representational organization. Within the present analyses, the principal fusiform signature of familiarity was increased representational correspondence, complementing the earlier temporal effect observed in LOC.

### Learning objectives reveal stage-dependent specialization

Manipulating model learning objective allowed us to ask which familiarity-sensitive dynamics are preferentially associated with face-identity learning and which also emerge under broader visual learning objectives. The FR and OR objectives provided contrasting supervisory targets, whereas the Dual objective involved object categorization with faces represented as one category rather than identity-level individuation. Long-term human familiarity cannot readily be reassigned experimentally, whereas models allow learning objectives to be compared within the same architecture and with the same loss function. This design therefore provides a within-architecture comparison of representational consequences associated with what the models were trained to recognize.

The LOC result provides the clearest indication that face-identity learning constrains, rather than uniquely determines, a familiarity-sensitive representation. Early familiar-face alignment was not exclusive to FR training: some architectures retained an early LOC solution under Dual or OR training. Across architectures, however, FR training made this early solution substantially more consistent, whereas broader objectives produced substantially more heterogeneous, architecture-dependent peak latencies. The relevant form of specialization therefore appears graded rather than all-or-none. Face-identity learning may not create an exclusive representational solution, but it appears to narrow the range of temporal solutions toward a more consistent early organization. This interpretation is consistent with computational evidence that face-identity optimization can impose specialized representational demands relative to object recognition (Abudarham et al., 2021), while task optimization more generally can shape functionally specialized representations without requiring completely segregated representational systems (Dobs et al., 2022, 2023). It is also compatible with evidence that category-selective representations can coexist with broader dimensions of object representation (Bao et al., 2020; Vinken et al., 2023).

The fusiform findings provide the complementary—and conceptually important—result. Familiar faces showed stronger fusiform correspondence than unfamiliar faces across all three learning objectives, and Dual- and OR-trained models showed greater peak correspondence for familiar faces than FR-trained models. The critical point is therefore not simply that fusiform cortex is familiarity-sensitive, but that this sensitivity is not uniquely associated with face-identity training. Models trained under broader visual objectives can therefor acquire representational structure that aligns with familiar-face neural representations, consistent with expertise and broader visual-learning accounts of visual recognition (Gauthier and Tarr, 2002) and with recent evidence that face-selective representations can coexist with broader object-coding dimensions (Bao et al., 2020; Vinken et al., 2023). Related computational work further shows that face-responsive representations can emerge in OR-trained networks without explicit face-recognition supervision (Xu et al., 2021). Together, these observations argue against equating functional specialization with representational exclusivity: face-relevant representational structure can emerge through broader visual learning, even if face-identity training imposes stronger constraints on other aspects of the representational hierarchy (Dobs et al., 2022, 2023).

The resulting picture is therefore neither strictly face-specific nor simply domain-general. Instead, specialization appears stage-dependent. Face-identity learning places particularly strong constraints on the temporal organization of intermediate LOC representations, whereas later familiar-face fusiform representations remain compatible with structure acquired through broader visual learning. This stage-dependent interpretation is conceptually consistent with recent primate evidence that face-selective inferotemporal populations can transition rapidly from a more general code supporting face detection to a face-specific code supporting fine face discrimination (Shi et al., 2026). Architecture dependence further qualifies this interpretation: under broader objectives, different architectures exhibited markedly different early and late LOC peak timings, whereas FR training narrowed the range of temporal solutions toward a more reproducible early organization. This is consistent with previous work showing that both network architecture and learning objective shape brain–model correspondence (Kubilius et al., 2019; Storrs et al., 2021; Prince et al., 2024). Recent work further shows that task-optimized models can predict training-induced representational plasticity in macaque inferior temporal cortex, providing convergent evidence that task optimization can capture aspects of experience-dependent representational change in high-level visual cortex (Sörensen et al., 2026).

The contrast with FaceNet illustrates why conclusions based on a single network can be misleading. Within FaceNet alone, the Dual objective produced particularly early and sustained LOC correspondence, which, considered in isolation, could suggest that Dual objective produces earlier familiar-face LOC alignment. Across architectures, however, the same objective produced substantially greater temporal heterogeneity. The stronger conclusion is therefore not that dual training generally produces earlier LOC alignment, but that FR training more consistently constrains familiar-face representations toward an early temporal solution, whereas broader objectives permit a wider and more architecture-dependent set of solutions.

### Scope, limitations, and future work

Brain–model alignment should be interpreted as correspondence between representational geometries, not as evidence that the brain and neural networks implement identical computations. More generally, task performance alone does not guarantee greater neural correspondence, and even high-performing visual networks can diverge from human and primate visual representations (Xu and Vaziri-Pashkam, 2021; Linsley et al., 2023). Several controls nevertheless constrain simpler explanations of the observed effects. Scrambled faces showed little higher-level correspondence despite measurable neural responses, trained networks consistently outperformed their untrained counterparts, and alignment was largely absent in the two nonvisual control regions. Together, these observations argue against explanations based solely on evoked-response magnitude, architecture alone, or spatially ubiquitous brain–model similarity.

Several limitations remain important. The MEG cohort was modest, source localization limits spatial precision, and peak-latency estimates can become unstable when similarity profiles are weak or multi-peaked. More generally, maximum-based summaries can be sensitive to the structure of the time × layer search space, motivating interpretation alongside the full time courses. The use of single presentations also imposed a signal-to-noise tradeoff: repeating unfamiliar identities could itself induce incidental familiarity, whereas avoiding repetition reduces the amount of within-subject data available for estimating representational geometry. In addition, the use of static face images limits generalization to dynamic and socially richer forms of face recognition. More generally, recent work on language-model–brain comparisons has shown that apparent brain–model alignment can depend on analysis choices and overlooked confounds, reinforcing the importance of conservative interpretation and strong controls (Hadidi et al., 2026). A further limitation concerns the level of inference in the primary model comparisons. Model-level analyses used subject-averaged MEG RDMs and treated architecture as the unit of comparison. These tests therefore assess consistency across the sampled models rather than population-level variability across human participants. Subject-level analyses provided complementary evidence, but the cross-objective conclusions are based primarily on the architecture-level comparisons.

The present behavioral task was designed primarily to maintain attention rather than measure face-recognition performance. Earlier LOC alignment should therefore be interpreted as earlier neural representational correspondence, not as evidence for faster behavioral recognition. Establishing whether the LOC timing and fusiform strength signatures predict recognition accuracy or response speed will require datasets in which behavioral performance is measured directly. An additional limitation concerns the computational manipulation itself. Although architecture and loss function were held constant across learning objectives within each architecture, and efforts were made to reduce gross differences in dataset scale and input resolution, the objectives necessarily relied on different image distributions. Learning objective is therefore not fully separable from visual training experience: differences across FR, OR, and Dual models may reflect the supervisory target, differences in the visual statistics of the training images, or both. In addition, the FR training corpus included 132 of the famous identities represented in the MEG stimulus set, with additional images collected where necessary. This overlap is relevant to the familiarity manipulation, but it limits what can be inferred from the greater cross-architecture consistency observed after FR training: the effect could reflect the demands of identity individuation, prior exposure to the specific identities, or both. A stronger future test would orthogonalize these factors by matching visual exposure while independently manipulating whether identities are individuated, categorized at the face-category level rather than individuated, or learned through broader object-recognition objectives (Yovel et al., 2023).

A particularly informative next step would be to track initially unfamiliar identities as familiarity develops. Such a longitudinal design could test whether learning progressively advances LOC alignment while separately strengthening later fusiform correspondence, providing a more direct test of the stage-dependent account. It would also allow these neural changes to be related to recognition accuracy and reaction time, addressing whether the representational signatures identified here have behavioral consequences. On the model side, extending the comparison to contrastive and self-supervised objectives would further test whether the observed effects depend specifically on supervised face recognition or reflect more general principles of representation learning. Recent work showing the emergence of category-selective representations under self-supervised objectives makes this an especially relevant direction (Prince et al., 2024), while brain-informed optimization provides a complementary route for testing how directly neural constraints shape learned representational geometry (Federer et al., 2020).

The LOC timing and fusiform correspondence signatures may also provide separable targets for studying impaired face recognition. Developmental prosopagnosia has been associated with atypical electrophysiological responses and alterations in occipitotemporal face-processing circuitry (Zhao et al., 2018; Manippa et al., 2023). Rather than predicting a unitary deficit, the present stage-dependent account motivates testing whether impaired recognition involves altered familiarity-related LOC timing, reduced familiar-face fusiform correspondence, or both. Model perturbations could help sharpen these alternatives by identifying which disruptions selectively affect each representational signature before testing these predictions in subject-level MEG data.

## Conclusion

Face familiarity affects successive stages of the ventral visual hierarchy in different ways. Face-identity learning most strongly constrains the temporal organization of intermediate LOC representations, whereas later familiar-face fusiform representations remain compatible with representational structure acquired through broader visual learning. Functional specialization in face recognition may therefore be better understood as graded and stage-dependent rather than all-or-none, with specialized constraints and broader learned representational structure contributing differently across the hierarchy. More broadly, combining millisecond-resolved MEG with systematic comparisons of model learning objective provides a framework for probing how different forms of visual learning relate to neural representational organization even when long-term experience cannot readily be manipulated in human observers.

## Methods

This section provides the necessary details to reproduce the work and results described in this article.

### MEG data

We analyzed publicly available magnetoencephalography (MEG) data from the multi-subject, multimodal face-perception dataset of Wakeman and Henson (2015). Data were acquired from 19 healthy participants (eight female; age range 23–37 years) using a 306-channel Elekta Neuromag Vectorview system. Three participants were excluded because of poor data quality, yielding a final sample of 16 participants. Participants viewed three stimulus types (*n* = 150 per condition): familiar faces depicting famous individuals, unfamiliar faces depicting non-famous individuals, and scrambled faces derived from the intact face stimuli. Familiar and unfamiliar face sets were approximately matched for age and sex (Fig. 1). All images were presented in grayscale and were cropped to show only the face. Stimuli were displayed for 800–1000 ms and each face was presented twice, with presentations either occurring immediately or following intervening stimuli. Only first presentations were considered for the present representational analyses to minimize repetition-related familiarity. MEG data were sampled at 1100 Hz. During acquisition, participants performed a symmetry-judgment task designed to maintain attention without explicitly requiring face recognition. Behavioral responses were therefore not used as a measure of familiarity-related recognition performance in the present study. Post-experiment familiarity debriefing confirmed the expected distinction between famous and non-famous faces at the group level.

MEG preprocessing and source reconstruction were implemented using MNE-Python version 1.6.1 (Gramfort et al., 2013) and followed procedures adapted from previous analyses of the same dataset (Jas et al., 2018).

#### Preprocessing

For each participant, the six MEG runs were transformed to the head position of the fourth run and processed using temporal signal-space separation implemented with MNE-Python’s `maxwell_filter`. Temporal signal-space separation used a 10-s window together with system-specific calibration and cross-talk compensation files. Bad MEG channels were identified from the existing MaxFilter logs for each run, and continuous head-position estimates were incorporated during Maxwell filtering. Following Maxwell filtering, MEG channels were low-pass filtered at 90 Hz using a zero-phase finite impulse response filter.

Events were aligned to stimulus presentation by correcting for the 34.5-ms delay between trigger onset and visual stimulus onset. Additional run-specific bad channels were interpolated before concatenating the six runs. Data were epoched from −200 to 800 ms relative to stimulus onset and baseline-corrected using the −200 to 0 ms prestimulus interval.

Independent component analysis (ICA) was performed using FastICA on the concatenated MEG recordings, retaining components explaining at least 99.9% of the variance. ICA was fitted to MEG channels after excluding marked bad channels. ECG-related components were identified using cross-trial phase statistics with a threshold of 0.8, and EOG-related components were identified from blink-related activity; at most three components of each type were removed. Automated amplitude rejection thresholds were subsequently estimated separately for each participant using the `autoreject` package and applied to remove residual artifactual epochs.

To ensure that brain and model representational dissimilarity matrices (RDMs) were constructed from a common stimulus set, stimuli unavailable for any participant or unavailable in the model stimulus set were excluded consistently across analyses. Only first-presentation trials from this common stimulus set were retained.

#### Source reconstruction

T1-weighted anatomical MRI images were processed using FreeSurfer’s `recon-all` (Fischl, 2012). Cortical source spaces were constructed on the reconstructed surfaces using oct6 spacing. A one-layer boundary-element model (BEM; conductivity = 0.3 S/m, ico = 4) was generated from the inner-skull surface, using watershed-derived BEM surfaces when necessary.

MEG and MRI coordinate systems were aligned using iterative closest-point coregistration. An initial six-iteration fit was followed by exclusion of head-shape points more than 5 mm from the scalp surface and a second 20-iteration fit. MEG forward solutions were computed using a minimum source-to-inner-skull distance of 5 mm.

Noise covariance matrices were estimated from the prestimulus interval using a shrunk covariance estimator with three-fold cross-validation. Minimum-norm inverse operators were constructed with a loose orientation constraint of 0.2 and depth weighting of 0.8. Source estimates were obtained using minimum-norm estimation with an assumed signal-to-noise ratio of 3 (*λ*^2^ = 1*/*9).

Participant-specific source estimates were morphed to the FreeSurfer `fsaverage` surface using an ico5 target source space. Source activity was then extracted from the 448 cortical parcels of the `aparc_sub` parcellation (Khan et al., 2018). For the representational analyses, activity across source vertices within each parcel was retained to characterize the spatial response pattern at each time point.

For complementary frequency-resolved analyses, morphed source estimates were band-pass filtered into theta (4–8 Hz), alpha (8–12 Hz), beta (12–30 Hz), gamma (30–55 Hz), and high-gamma (55–90 Hz) bands. The analytic amplitude envelope was then obtained using the Hilbert transform and processed using the same parcel-based framework.

### Artificial Neural Networks

#### Model architectures

We compared seven convolutional neural network architectures spanning a range of architectural designs: ResNet-50 (He et al., 2016), VGG-16 with batch normalization (Simonyan and Zisserman, 2015), Inception-v3 (Szegedy et al., 2016), MobileNetV2 (Sandler et al., 2018), CORnet-S (Kubilius et al., 2019), FaceNet, implemented with an Inception-ResNet-v1 backbone (Schroff et al., 2015), and SphereFace (Liu et al., 2017). The first five architectures were originally developed primarily for object recognition or general visual classification, whereas FaceNet and SphereFace were developed for face recognition. All models were implemented in PyTorch (Paszke et al., 2019).

To enable comparison of learning objectives while holding architecture-specific training losses constant, separate instances of the same architectural backbone were used across objectives and all models were optimized using cross-entropy classification loss. Thus, the specialized metric-learning or angular-margin losses associated with the original FaceNet and SphereFace formulations were not used in the present comparisons. For each architecture, a randomly initialized untrained counterpart was also included to assess the contribution of learned weights to brain–model correspondence beyond architecture alone. Learning objective was treated as the primary model-side manipulation, whereas variation across architectures was used to assess the stability and generalization of the observed effects across different architectural inductive biases.

#### Training datasets and learning objectives

Here, *learning objective* refers to what a network was trained to recognize or discriminate, rather than to the mathematical loss function used for optimization. Separate instances of each architecture were optimized under three learning objectives: face-identity recognition (FR), object-category recognition (OR), and object categorization including a face category (Dual) (Fig. 1).

For the FR objective, models were first trained for face-identity classification on VGGFace2 (Cao et al., 2018) and subsequently fine-tuned on a CelebA-derived identity dataset (Liu et al., 2015). The CelebA training set comprised approximately 1000 identities and was constructed to include 132 of the famous identities represented in the MEG experiment. When CelebA contained insufficient images for a target identity, additional photographs of the same individual were collected from the web. Face images were cropped using facial landmark detection, and the identity classes were balanced to contain approximately 30 images per identity. The supervisory target in this condition was the identity of the individual depicted in each image.

For the OR objective, models were trained to classify object categories using a subset of ImageNet (Deng et al., 2009). For the Dual objective, the same object-categorization framework was augmented with a face category containing images from multiple identities drawn from the CelebA distribution. Importantly, these faces were assigned a common face-category label rather than separate identity labels. The Dual objective therefore required distinguishing faces from other object categories but did not require identity individuation.

To reduce differences unrelated to learning objective, ImageNet was subsampled and the training sets were constructed to approximate the scale of the face-training datasets, while all model inputs were standardized to a spatial resolution of 224 *×* 224 pixels. These controls reduced gross differences in training exposure and input resolution across objectives. However, the datasets necessarily retained different image statistics and class structures; learning objective therefore cannot be completely separated from visual training experience, as considered in the limitations.

For the face datasets, images were partitioned within identity into approximately 70% training, 15% validation, and 15% test sets. For ImageNet, the selected training subset was used for model fitting, and the available validation data were divided into validation and held-out test subsets.

#### Network training

Within each architecture, the same architectural backbone and cross-entropy classification loss were used across learning objectives. Hyperparameter optimization involved a random search across tasks and architectures. Models were optimized using stochastic gradient descent with momentum (0.9) and weight decay (10^−4^). The initial learning rate was 0.01 for FR training and 0.001 for OR and Dual training, and was adjusted using a StepLR scheduler every 10 epochs. Models were trained for 50 epochs, with training and validation loss and accuracy monitored throughout training using Weights & Biases (WandB). Performance was subsequently evaluated on held-out test data.

Random seeds were fixed across Python, NumPy, and PyTorch, and deterministic cuDNN settings were enabled to improve reproducibility. Because optimization settings differed where necessary to obtain stable learning across objectives, the training conditions should not be interpreted as differing solely in supervisory target. The principal controls were instead the use of the same architectural backbone and classification loss within each architecture, together with efforts to match dataset scale and input resolution.

The purpose of model optimization was not to establish state-of-the-art performance on the respective computer-vision benchmarks, but to obtain learned representations under contrasting recognition objectives while preserving comparable architectural and training conditions. Model weights were retained after training for subsequent representational analyses.

#### Model activations

To construct model representations for comparison with the MEG data, the face stimuli from the MEG experiment were presented to each trained and untrained network. Activations were extracted from successive hidden layers using forward hooks and flattened into feature vectors while preserving the stimulus dimension. These activation patterns were subsequently used to construct layer-specific representational dissimilarity matrices. The same common set of 428 stimuli retained for the MEG analyses was used for the final brain–model comparisons.

### Representational Similarity Analysis

We used Representational Similarity Analysis (RSA) (Kriegeskorte et al., 2008) to quantify correspondence between the representational geometry of source-resolved MEG activity and that of the convolutional neural networks. RSA tests whether stimuli that are represented similarly in one system are also represented similarly in another by comparing their representational dissimilarity matrices (RDMs).

#### Representational Dissimilarity Matrices

For both MEG and network representations, pairwise dissimilarity between stimuli was quantified using Pearson correlation distance (1 − *r*). For the common set of 428 stimuli retained after preprocessing, this yielded a 428 *×* 428 RDM, with values ranging from 0 for perfectly correlated response patterns to 2 for perfectly anticorrelated response patterns. Because RDMs are symmetric, only the upper triangular elements excluding the diagonal were used in subsequent analyses. Condition-specific sub-RDMs were then extracted for familiar faces (130 stimuli), unfamiliar faces (150 stimuli), and scrambled faces (148 stimuli) (Fig. 1).

### Brain RDMs

For each participant, cortical parcel, and time point, each stimulus was represented by the pattern of source activity across the vertices contained within that parcel. For a parcel containing *N* source vertices, this yielded one *N*-dimensional response vector per stimulus. Pearson correlation distance between all pairs of stimulus-response vectors was then used to construct a time-resolved RDM for each of the 448 cortical parcels.

The same procedure was applied to the frequency-resolved source estimates, replacing broadband source activity with the Hilbert amplitude envelope within each frequency band. RDMs were computed independently for each participant. For the primary model-level analyses, the resulting RDMs were averaged element-wise across the 16 participants before comparison with the network representations. Thus, participant averaging was performed at the level of representational geometry rather than on the source signals themselves. Complementary subject-level analyses retained individual-participant RDMs to assess the robustness of the principal effects across participants.

### Network RDMs

We characterized representations from successive hidden layers of each of the seven architectures under the FR, OR, Dual, and untrained conditions. The MEG stimulus images were presented to each network, and layer activations were captured using registered forward hooks. Activation tensors were detached from the computation graph and flattened into feature vectors while preserving the stimulus dimension. Pairwise Pearson correlation distance (1 − *r*) between these vectors was used to construct a layer-specific RDM. Condition-specific familiar, unfamiliar, and scrambled sub-RDMs were extracted using the same stimulus subsets as for the MEG data.

### Brain–model similarity

Brain–model correspondence was quantified by comparing the representational geometry of each MEG RDM with that of each network layer (Fig. 1h). Specifically, the upper triangular elements of the corresponding MEG and network RDMs were vectorized and compared using Spearman rank correlation. The resulting RSA coefficient therefore quantified the correspondence between the relative pairwise representational structure of the stimuli in the brain and model representations.

This procedure was repeated across cortical parcels, time points, network layers, stimulus conditions, architectures, and learning objectives, yielding time- and layer-resolved maps of brain–model representational correspondence. The same analysis was applied to broadband MEG activity and to each of the five frequency-resolved amplitude envelopes. Unless otherwise stated, the primary model comparisons used the subject-averaged MEG RDMs described above, whereas subject-level RSA was used as a complementary robustness analysis.

For analyses of alignment strength, peak similarity was defined as the maximum brain–model RSA value across the time *×* layer space for a given region, model, and stimulus condition. For analyses of temporal organization, peak latency was defined as the time coordinate of maximal similarity within the 50–400 ms post-stimulus window. Peak latency therefore indexes when correspondence between the neural and model representational geometries was maximal; it should not be interpreted as the latency of a univariate neural response, the speed of neural computation, or behavioral recognition. Because maximum-based estimates can be sensitive to weak or multi-peaked similarity profiles, peak-latency comparisons were interpreted together with the complete time-resolved similarity profiles rather than in isolation.

### Similarity scores and noise ceiling

A noise ceiling was estimated separately for each cortical parcel, time point, stimulus condition, and signal type to quantify the maximum representational correspondence expected given the reliability of the MEG data (Nili et al., 2014). We used the upper noise ceiling, computed by correlating each participant’s RDM with the grand-average RDM across all participants and then averaging these correlations across participants. Noise-ceiling estimation used the same Spearman correlation measure as the brain–model RSA.

To account for variation in the reliability of MEG representational geometry across regions, time points, and stimulus conditions, each brain–model RSA coefficient was normalized by its corresponding upper noise-ceiling estimate. All similarity values reported in the principal analyses therefore represent noise-ceiling-normalized brain–model correspondence.

### Statistical testing

#### Regions of interest and event-related responses

Primary analyses focused on three regions along the ventral visual pathway: right pericalcarine cortex (V1), right lateral occipital cortex (LOC), and right fusiform cortex. These regions were motivated by their successive positions in the visual hierarchy and by previous work implicating occipital and fusiform cortex in face perception and recognition. Importantly, region selection was performed without reference to the CNN–MEG RSA results.

As a complementary characterization of the source-localized MEG responses, event-related fields (ERFs) were computed by averaging epochs within each stimulus condition. Condition effects were examined for familiar versus scrambled faces, unfamiliar versus scrambled faces, and familiar versus unfamiliar faces. Statistical significance across time was assessed using cluster-based one-sample permutation tests implemented in MNE-Python with 1000 permutations. Clusters with *p <* 0.05 were considered significant. These response-level analyses were used to characterize condition-sensitive neural activity and were evaluated separately from the representational similarity analyses.

#### Time-resolved brain–model correspondence

Statistical significance of the time-resolved RSA maps was assessed using cluster-based permutation tests. Multiple comparisons were controlled at the cluster level across network layers, regions, and time, with the family-wise error rate set to *α* = 0.05. In the figures, significant clusters are displayed when they span at least 10 consecutive time samples. The same procedure was applied to the broadband and frequency-resolved RSA analyses. Because cluster boundaries depend on the temporal extent and adjacency of the effect, significant clusters were interpreted as evidence for sustained periods of brain–model correspondence rather than as precise estimates of effect onset or offset.

#### Peak similarity and peak-latency comparisons

The primary comparisons of peak similarity and peak latency were performed at the model level. As described above, these analyses used MEG RDMs averaged across the 16 participants and treated the seven network architectures as the inferential units (*n* = 7). Within-architecture comparisons between stimulus conditions, learning objectives, or cortical regions were therefore evaluated using two-sided paired *t*-tests across architectures (*df* = 6). Pairing by architecture controls for baseline differences in correspondence attributable to architectural design when testing the effects of familiarity or learning objective.

Peak-similarity analyses compared the maximum noise-ceiling-normalized RSA value across the time *×* layer space for each architecture and condition. Peak-latency analyses compared the corresponding time of maximal similarity within the predefined 50–400 ms post-stimulus window. Because peak estimates summarize a larger time *×* layer search space and can be sensitive to weak or multi-peaked similarity profiles, inferential conclusions from these measures were interpreted together with the corresponding time-resolved RSA profiles.

The full pairwise-comparison matrices shown in the figures provide a descriptive overview of differences among stimulus conditions and learning objectives. Inferential conclusions in the text were restricted to contrasts directly addressing the study questions, namely familiarity effects within learning objectives and differences among learning objectives for a given stimulus condition.

#### Subject-level robustness analyses

To complement the model-level analyses, selected effects were evaluated at the participant level by correlating each participant’s MEG RDMs separately with the corresponding network RDMs. For these analyses, participants constituted the inferential units (*n* = 16), and within-participant differences between conditions were assessed using two-sided paired *t*-tests (*df* = 15). These analyses were treated as robustness tests of the regional patterns identified in the primary subject-averaged model comparisons rather than as the basis for inference about differences among learning objectives across architectures.

Unless otherwise stated, statistical tests were two-sided and used an alpha level of 0.05.

## Data and Code

The MEG data analyzed in this study are publicly available through OpenNeuro under accession `ds000117` (https://openneuro.org/datasets/ds000117) (Wakeman and Henson, 2015). The face- and object-image datasets used for model training (CelebA, VGGFace2, and ImageNet) are available from their respective original sources, as cited above.

All code used for MEG preprocessing and source reconstruction, neural-network training and activation extraction, RDM construction, representational similarity analysis, and generation of the reported results is publicly available at https://github.com/BabaSanfour/CNN-MEG-FaceProcessing. The repository also contains the scripts used to construct the training-data subsets and reproduce the analysis pipeline.

The trained model weights used in the analyses will be made publicly available upon publication.

## Acknowledgments

The authors acknowledge the Digital Research Alliance of Canada and Calcul Québec for providing advanced research computing resources used in this work. The authors also thank Wakeman and Henson for making the multimodal face-perception dataset used in this study publicly available.

## Funding

H.A. was supported by scholarships from Cerebrum, the Centre interdisciplinaire de recherche sur le cerveau et l’apprentissage (CIRCA), and the Centre UNIQUE (Unifying Neuroscience and Artificial Intelligence in Quebec). H.A. is currently supported by a Merit Scholarship from the AI Research Fund of the Faculty of Medicine, Université de Montréal. Travel to present this work was supported by the Centre UNIQUE, the Computational and Systems Neuroscience (COSYNE) conference, and the Réseau de Bio-Imagerie du Québec (RBIQ). S.B. is funded by NSERC Discover Grant (RGPIN-2023-03875). K.J. is supported by funding from the Canada Research Chairs (950-232368) program and a Discovery Grant from the Natural Sciences and Engineering Research Council of Canada (2021-03426).

## Supplementary Information

**Figure S1.**
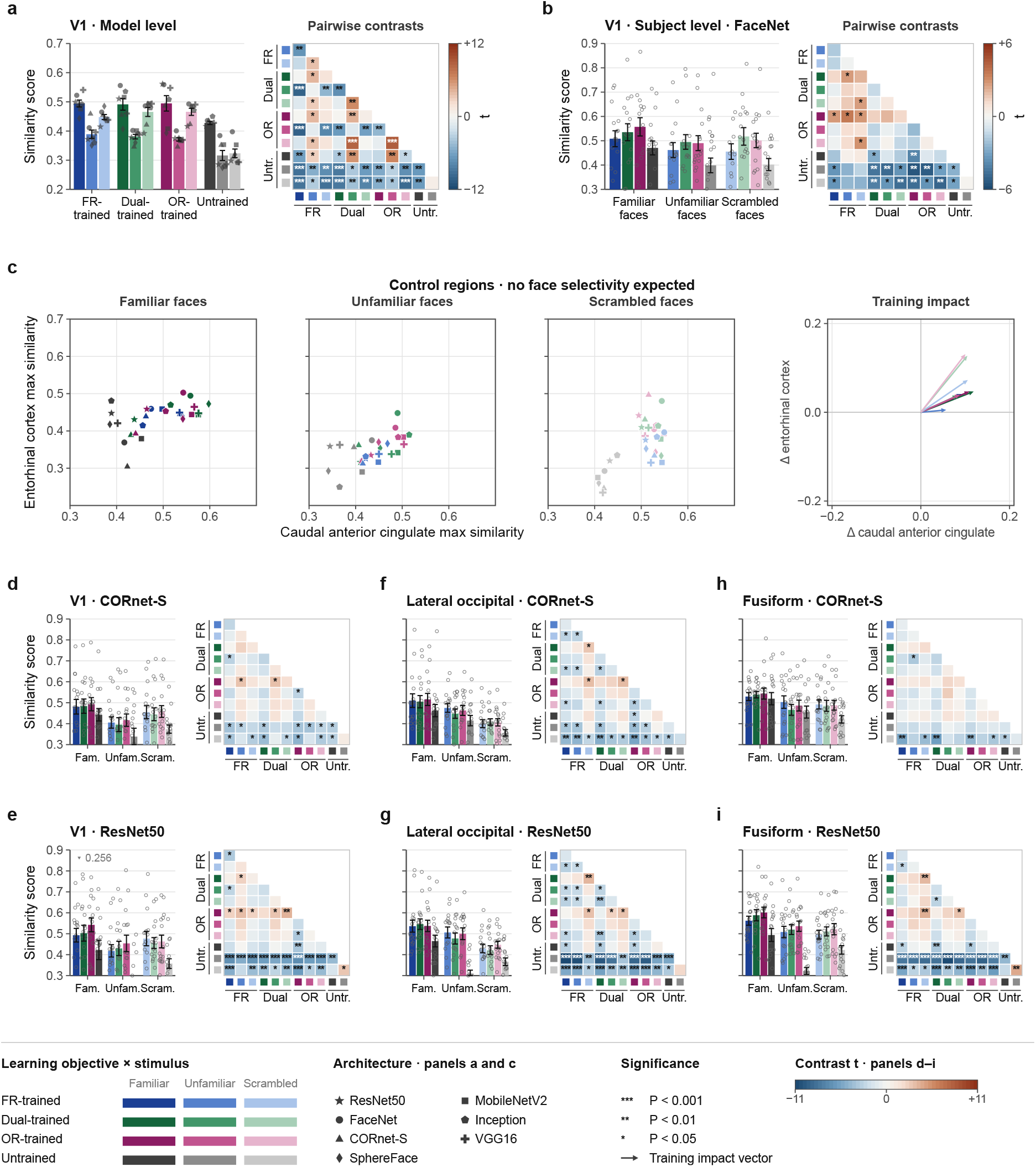
MEG–CNN similarity in V1 and control regions, with subject-level robustness across architectures. (**a**) Model-level peak MEG–CNN RSA similarity in V1, defined as the maximum similarity across time and network layers. Similarity was computed from subject-averaged MEG RDMs (*n*=16) for seven CNN architectures (*n*=7; each point represents one architecture) across learning objectives and stimulus conditions. The adjacent heat map shows pairwise contrasts, expressed as *t* values from two-sided paired tests across architectures. (**b**) Subject-level FaceNet analysis in V1. Bars show peak similarity for familiar, unfamiliar, and scrambled faces across participants (*n*=16), with individual participants shown as points. The adjacent heat map summarizes pairwise contrasts with *t* values from two-sided paired tests across participants. (**c**) Control-region analysis comparing peak similarity in entorhinal cortex and caudal anterior cingulate cortex, shown separately for familiar, unfamiliar, and scrambled faces. Each point represents one architecture. Training-impact vectors show the change from each architecture’s untrained baseline to its trained counterpart. (**d–i**) Subject-level robustness analyses for two additional architectures. CORnet-S results are shown for V1 (d), lateral occipital cortex (LOC; f), and fusiform cortex (h); ResNet50 results are shown for V1 (e), LOC (g), and fusiform cortex (i). Each point represents one participant, and adjacent heat maps summarize pairwise contrasts across learning objective *×* stimulus conditions using two-sided paired tests across participants.

**Figure S2.**
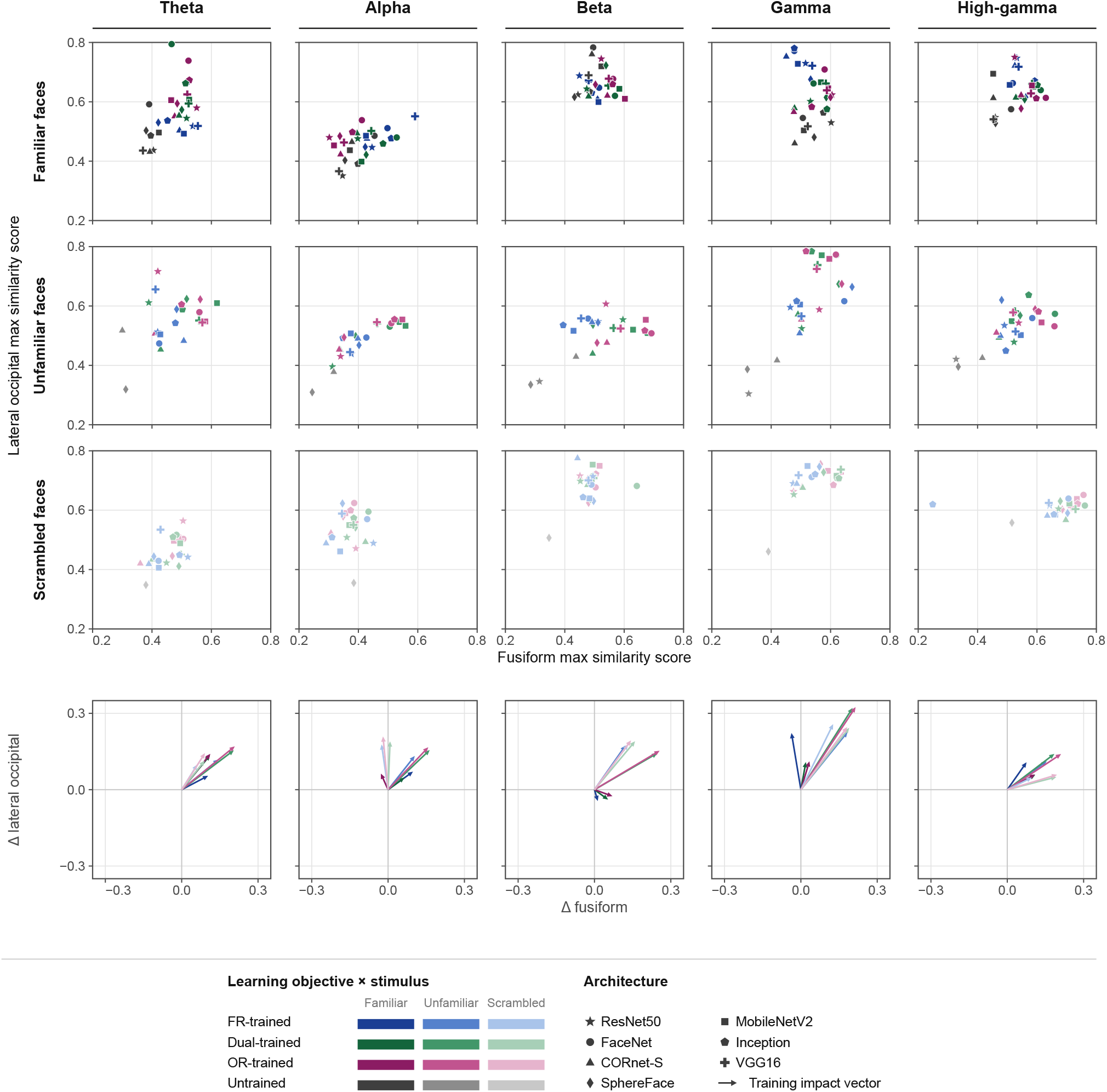
Frequency-resolved MEG–CNN similarity in lateral occipital and fusiform cortex. Peak MEG–CNN RSA similarity is shown separately for theta, alpha, beta, gamma, and high-gamma activity. The upper three rows compare lateral occipital cortex (LOC) and fusiform peak similarity for familiar, unfamiliar, and scrambled faces, respectively. Peak similarity was defined as the maximum across time and network layers within each frequency band; each point represents one of seven CNN architectures under a given learning objective and stimulus condition. The bottom row shows training-dependent changes in LOC and fusiform similarity, with vectors connecting each architecture’s untrained baseline to its trained counterpart.

**Figure S3.**
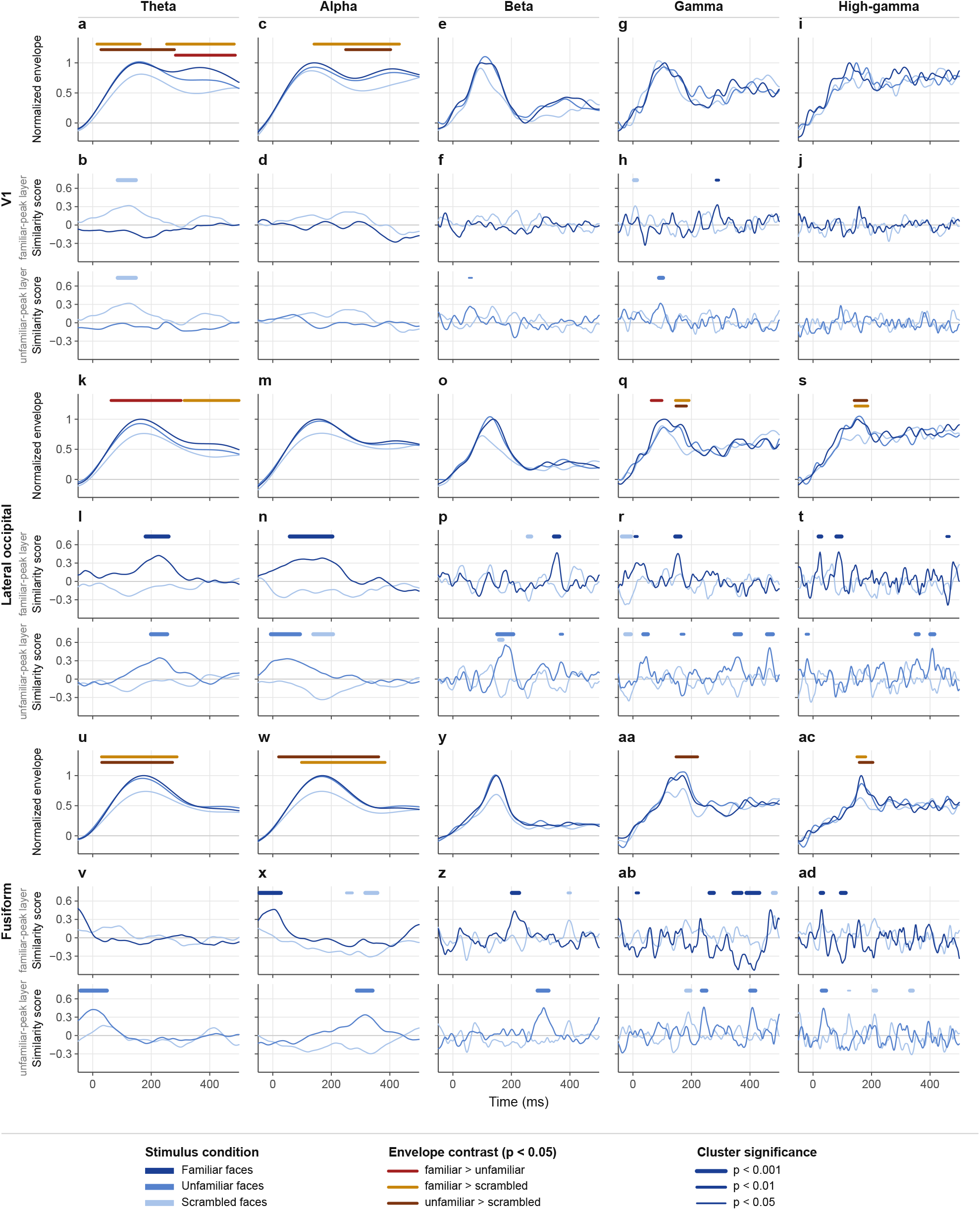
Frequency-resolved MEG–FaceNet similarity and band-limited source responses across V1, lateral occipital cortex, and fusiform cortex. (**a–j**) V1, (**k–t**) lateral occipital cortex (LOC), and (**u–ad**) fusiform cortex, shown separately for theta, alpha, beta, gamma, and high-gamma activity. For each frequency band and region, the upper panel shows the normalized source-level amplitude envelope for familiar, unfamiliar, and scrambled faces. Colored bars indicate significant condition differences in the MEG response (*p <* 0.05; yellow = familiar *>* scrambled, orange = unfamiliar *>* scrambled, red = familiar *>* unfamiliar). The two lower panels show time-resolved MEG–FaceNet RSA similarity at the FaceNet layer yielding maximal similarity for familiar faces and unfamiliar faces, respectively. Subject-averaged, time-resolved MEG RDMs were correlated with layer-resolved FaceNet RDMs at each time point. Horizontal blue bars indicate significant above-chance similarity clusters, assessed with cluster-based permutation tests and family-wise correction across layers, regions, and time; clusters are displayed when spanning at least 10 consecutive samples (*p <* 0.05).

**Figure S4.**
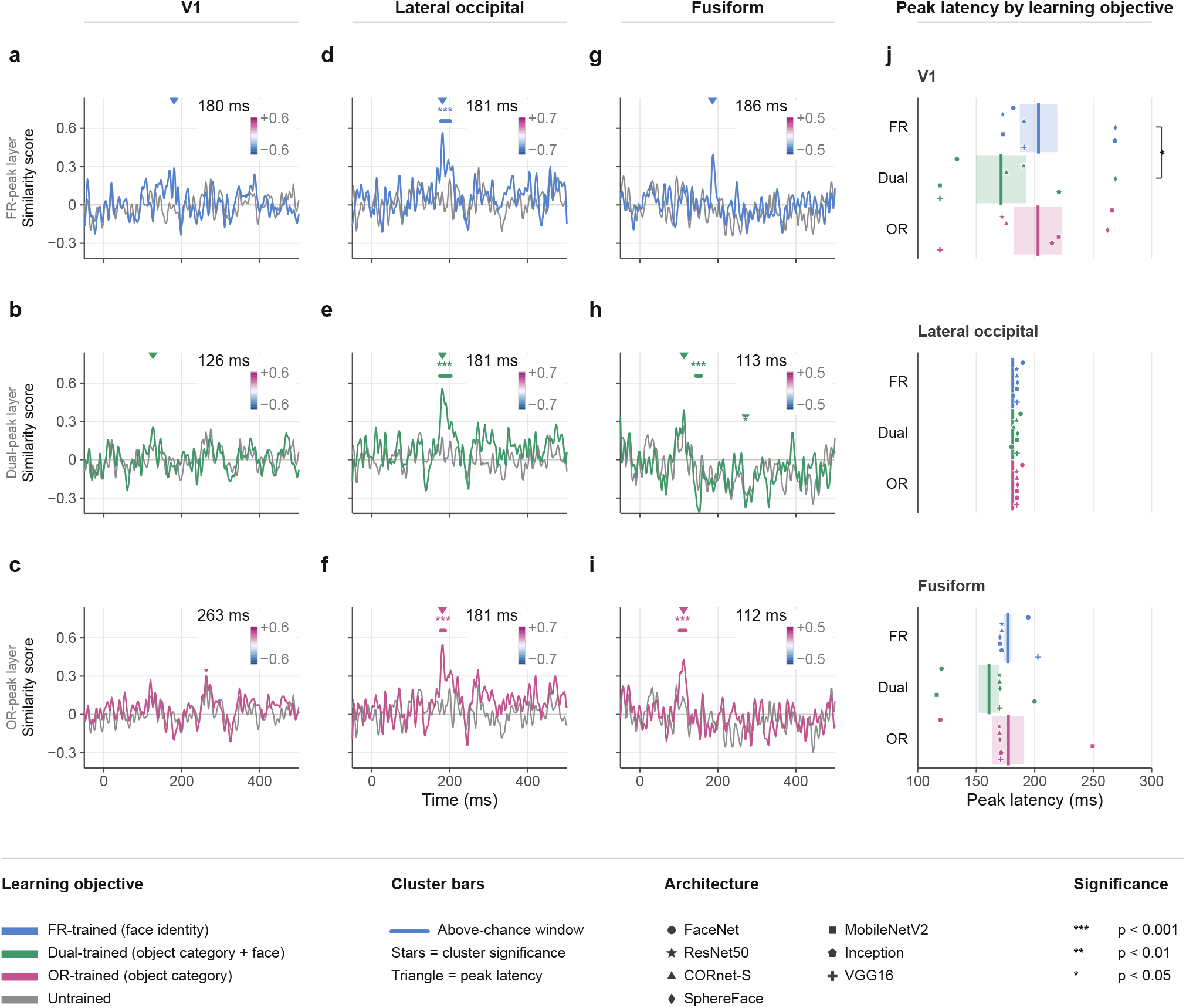
Learning objective shapes the spatiotemporal dynamics of MEG–CNN similarity for unfamiliar faces across V1, lateral occipital cortex, and fusiform cortex. (**a**,**d**,**g**) Time-resolved MEG–FaceNet RSA similarity for unfamiliar faces under the FR-trained objective in V1 (a), lateral occipital cortex (LOC; d), and fusiform cortex (g). Similarity is shown at the CNN layer yielding maximal unfamiliar-face similarity for the FR-trained model; gray traces show the corresponding untrained model. Colored horizontal bars indicate significant above-chance similarity clusters, with stars denoting significance level, and triangles marking peak similarity latency. Inset cortical maps show the spatial distribution of similarity at the corresponding peak latency. (**b**,**e**,**h**) Corresponding analyses for the Dual-trained objective in V1 (b), LOC (e), and fusiform cortex (h), evaluated at the layer yielding maximal unfamiliar-face similarity for the Dual-trained model. (**c**,**f**,**i**) Corresponding analyses for the OR-trained objective in V1 (c), LOC (f), and fusiform cortex (i), evaluated at the layer yielding maximal unfamiliar-face similarity for the OR-trained model. (**j**) Peak latency of MEG–CNN alignment for unfamiliar faces across V1, LOC, and fusiform cortex, shown separately for FR-trained, Dual-trained, and OR-trained models. Peak latency was defined as the time of maximal similarity within the 50–400 ms search window. Each point represents one of seven CNN architectures; bars indicate mean *±* SD. Brackets denote significant paired comparisons across regions and learning objectives.

## Notes

### Competing Interest Statement

The authors have declared no competing interest.

